# A small proportion of Talin molecules transmit forces to achieve muscle attachment *in vivo*

**DOI:** 10.1101/446336

**Authors:** Sandra B. Lemke, Thomas Weidemann, Anna-Lena Cost, Carsten Grashoff, Frank Schnorrer

## Abstract

Cells in a developing organism are subjected to particular mechanical forces, which shape tissues and instruct cell fate decisions. How these forces are sensed and transmitted at the molecular level is thus an important question, which has mainly been investigated in cultured cells *in vitro*. Here, we elucidate how mechanical forces are transmitted in an intact organism. We studied *Drosophila* muscle attachment sites, which experience high mechanical forces during development and require integrin-mediated adhesion for stable attachment to tendons. Hence, we quantified molecular forces across the essential integrin-binding protein Talin, which links integrin to the actin cytoskeleton. Generating flies expressing three FRET-based Talin tension sensors reporting different force levels between 1 and 11 pN enabled us to quantify physiologically-relevant, molecular forces. By measuring primary *Drosophila* muscle cells, we demonstrate that *Drosophila* Talin experiences mechanical forces in cell culture that are similar to those previously reported for Talin in mammalian cell lines. However, *in vivo* force measurements at developing flight muscle attachment sites revealed that average forces across Talin are comparatively low and decrease even further while attachments mature and tissue-level tension increases. Concomitantly, Talin concentration at attachment sites increases five-fold as quantified by fluorescence correlation spectroscopy, suggesting that only few Talin molecules are mechanically engaged at any given time. We therefore propose that high tissue forces are shared amongst a large excess of adhesion molecules of which less than 15% are experiencing detectable forces at the same time. Our findings define an important new concept of how cells can adapt to changes in tissue mechanics to prevent mechanical failure *in vivo*.

## Introduction

The shape of multicellular organisms critically depends on the presence of mechanical forces, during development [1,2]. Forces not only generate form and flows within tissues [3,4] but can also control cell fate decisions [5,6] or trigger mitosis [7]. There are various ways to quantify forces at the cellular or tissue level [8,9], however mechanical forces experienced by proteins in cells have only recently become quantifiable with the development of Förster Resonance Energy Transfer (FRET)-based molecular tension sensors [10]. These sensors contain a donor and an acceptor fluorophore connected by a mechano-sensitive linker peptide, which reversibly unfold and extend when experiencing mechanical forces. As a result, such sensors report forces as a decrease in FRET efficiency resulting from an increase in distance between the fluorophores. Since previous studies analysed molecular forces using *in vitro* cell culture systems [11-16] and insights from *in vivo* experiments are still limited [17-20], it remains largely open how mechanical loads are processed at the molecular level in tissues of living organisms.

Integrins are a major and highly conserved force bearing protein family. They connect the actomyosin cytoskeleton to the extracellular matrix and are essential for numerous mechanically regulated processes *in vivo* or *in vitro* [21,22]. However, *in vivo* it is particularly unclear how integrin-based structures are mechanically loaded since forces have so far been analysed in focal adhesions, which typically are not found in soft tissues [11-13,16]. Therefore, we chose to investigate *Drosophila* muscle attachment sites *in vivo*, which experience high mechanical forces during development [23] and depend on integrin-based attachment of muscle fibers to tendons cells [21,24]. For the molecular force measurements we selected the integrin activator and mechanotransducer Talin, which is essential for all integrin mediated functions and binds with its globular head-domain to the tail of β-integrin and with its rod-domain to actin filaments [25,26]. Thus, Talin is in the perfect position to sense mechanical forces across integrin-dependent adhesive structures. Surprisingly, we find that less than 15% of the Talin molecules experience significant forces at muscle attachments *in vivo* suggesting that high tissue forces are rather sustained by recruiting a large excess of Talin molecules to muscle attachments. This may have important impact for the robustness of muscle attachment under peak mechanical load in muscles.

## Results

### A *Drosophila* Talin tension sensor

To enable quantitative force measurements, we generated various *Drosophila* Talin tension sensor and control flies by modifying the endogenous *talin* (*rhea*) gene using a two-step strategy based on CRISPR/Cas9 genome engineering and ϕC31-mediated cassette exchange (Fig. 1a, Extended data Fig. 1) [27]. This strategy enabled us to generate an entire set of Talin tension sensor fly lines with YPet and mCherry (mCh) FRET pairs and three different mechano-sensitive linker peptides [11,13], Flagelliform (F40), Villin headpiece (HP) and its stable variant (HPst), reporting forces of 1-6 pN, 6-8 pN and 9-11 pN, respectively (Fig. 1b). The sensor modules were inserted both internally between the Talin head- and rod-domains (F40-TS, TS, stTS) at the analogous position used in mammalian Talin to report forces *in vitro* [11,16], and C-terminally as a zero-force control (C-F40-TS, C-TS, C-stTS). Furthermore, the individual fluorescent proteins were inserted at both positions as controls (I-YPet, I-mCh, C-YPet, C-mCh). Importantly, all stocks are homozygous viable, fertile and do not display any overt phenotype indicating that the Talin tension sensor proteins are functional.

**Fig. 1.**
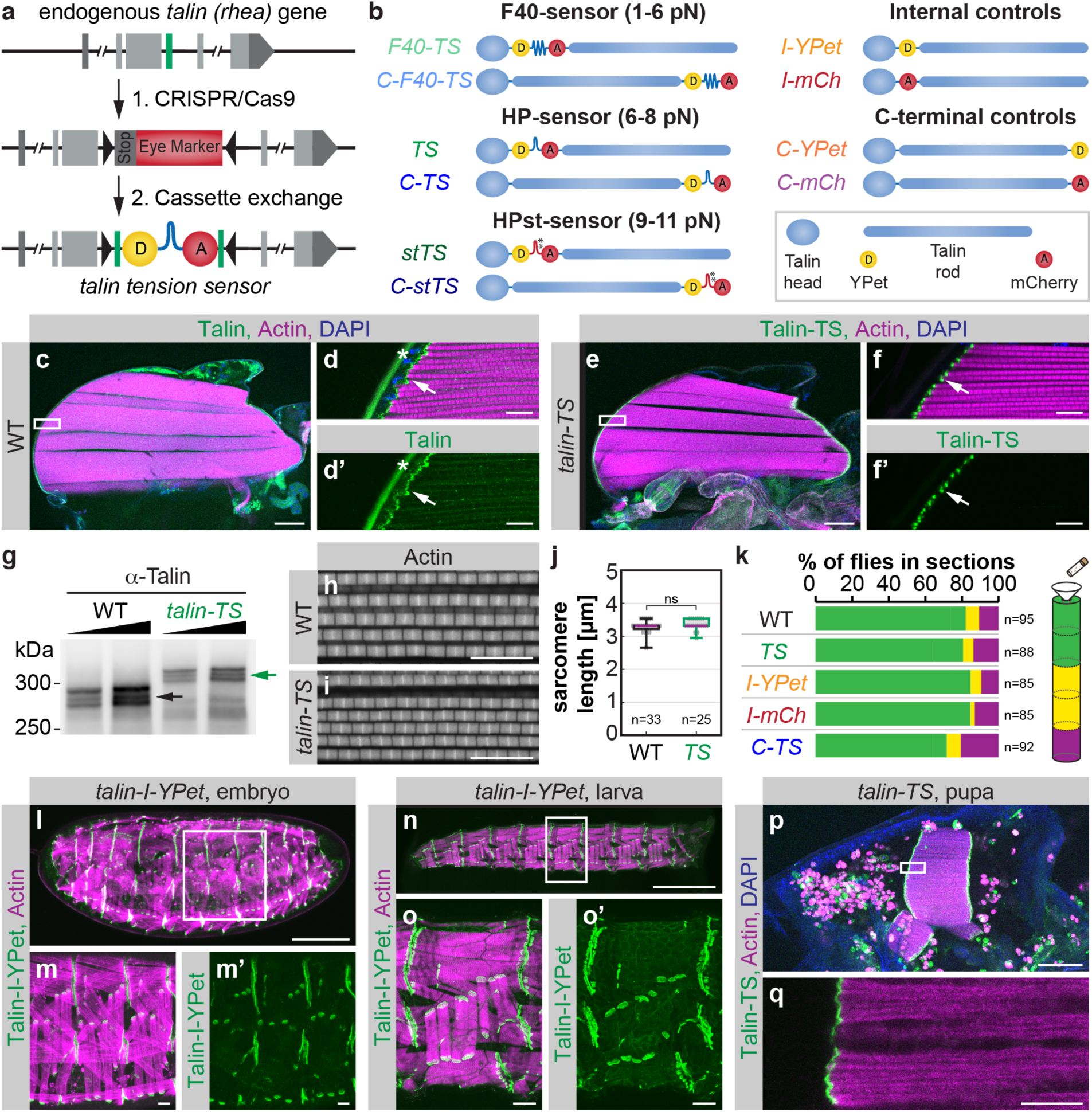
Talin tension sensor generation and verification. **a**, 2-step genome engineering strategy of the *talin* (*rhea*) gene. Step 1: Cas9-mediated insertion of an eye marker cassette replacing the target exon (green). Step 2: ϕC31-mediated cassette exchange restoring the original exon and including a tension sensor. See Extended Data Fig. 1 for details. **b**, Overview of Talin tension sensor and control flies. Sensors with three different mechano-sensitive linker peptides, F40, HP and HPst, were generated. Respective force regimes are indicated. Each sensor was inserted internally (F40-TS, TS, stTS) or at the C-terminus (C-F40-TS, C-TS, C-stTS). Individual fluorescent protein controls were also generated (I-YPet, I-mCh, C-YPet, C-mCh). **c-d**, Wild-type (WT) adult hemithorax stained with Talin antibody, phalloidin (Actin) and DAPI. White box in **c** indicates zoom-in area shown in **d** and **d’**. Note the Talin localization at myofibril tips (arrow). The star indicates background fluorescence from the cuticle. **e-f**, *talin tension sensor* (*talin-TS*) adult hemithorax showing Talin-TS localization at myofibril tips (arrow). **g**, Western blot of whole fly extract from WT and *talin-TS* flies probed with Talin antibody. Note the up-shift of all Talin-TS bands (green arrow) compared to WT (black arrow). **h-j**, Phalloidin stainings of adult hemithoraxes showing normal sarcomere morphology in WT (**h**) and *talin-TS* (**i**) flies and normal sarcomere length (**j**) (Mann Whitney test, ns=not significant, p=0.40, n=number of flies). **k**, Flight test (two-way ANOVA, no significant differences compared to WT in 6 replicates, n=total numbers of flies). **l-q**, Talin-I-YPet or Talin-TS expression at different stages of development. Live images of a stage 17 *Talin-I-YPet* embryo (**l-m**) and an L3 larva (**n-o**) co-expressing *Mef2-GAL4, UAS-mCherry-Gma* as a muscle actin marker. (Since the actin marker contains mCherry we used Talin-I-YPet here). A 32 h APF *talin-TS* pupa (**p-q**) stained with phalloidin and DAPI. Scale bars are 100 μm in **c, e, l, o** and **p**, 10 μm in **d, f, m** and **q**, and 1 mm in **n**.

To assess the functionality of the Talin tension sensor protein (Talin-TS) more rigorously, we first analysed Talin-TS localization in adult hemi-thoraxes and found that Talin-TS localizes to myofibril tips as expected (Fig. 1c-f). Second, we performed western blot analysis to ensure that the tension sensor module is incorporated into Talin protein isoforms as designed (Fig. 1g). Third, we quantified sarcomere length in flight muscles and found the expected length of 3.2 μm in wild type (WT) [28] and *talin-TS* flies (Fig. 1h-j). Forth, we tested flight ability [29] and found that the insertion of neither the sensor module or the individual fluorescent proteins into the internal position nor the sensor module at the C-terminus caused flight defects (Fig. 1k). Finally, we confirmed that Talin-TS (or Talin-I-YPet) is expressed correctly at all developmental stages (embryo, larva and pupa) and is detected most prominently at muscle attachment sites as previously reported for endogenous Talin (Fig. 1l-q) [30]. Together, these data demonstrate that the tension sensor module is properly incorporated into Talin and the resulting protein is functional. This permits the quantification of mechanical tension across Talin in any tissue and at any developmental stage of *Drosophila in vivo*.

### Forces across *Drosophila* Talin in primary muscle fiber cultures

To ensure that our approach is comparable to previous Talin force measurements in cultured mammalian cells, we established muscle fiber cultures by incubating primary myoblasts *in vitro* for 5-7 days [31,32]. Isolated myoblasts from *talin-I-YPet* embryos differentiated into striated, often multinucleated muscle fibers and efficiently adhered to the underlying plastic substrate (Fig. 2a, b). In these cells, Talin-I-YPet localises to adhesions at the fiber tips and at myofibril ends as well as to costameres, which connect myofibrils at the sarcomeric Z-discs to the cell membrane [33]. Primary muscle fibers generated from *talin-I-YPet*, *talin-TS* and *talin-C-TS* embryos display similar morphologies (Fig. 2c-e) and contract spontaneously (Supplementary Video 1). Adhesions at the fiber tips do not move during these contractions while costameres are mobile and thus not fixed to the plastic substrate (Fig. 2f).

**Fig. 2.**
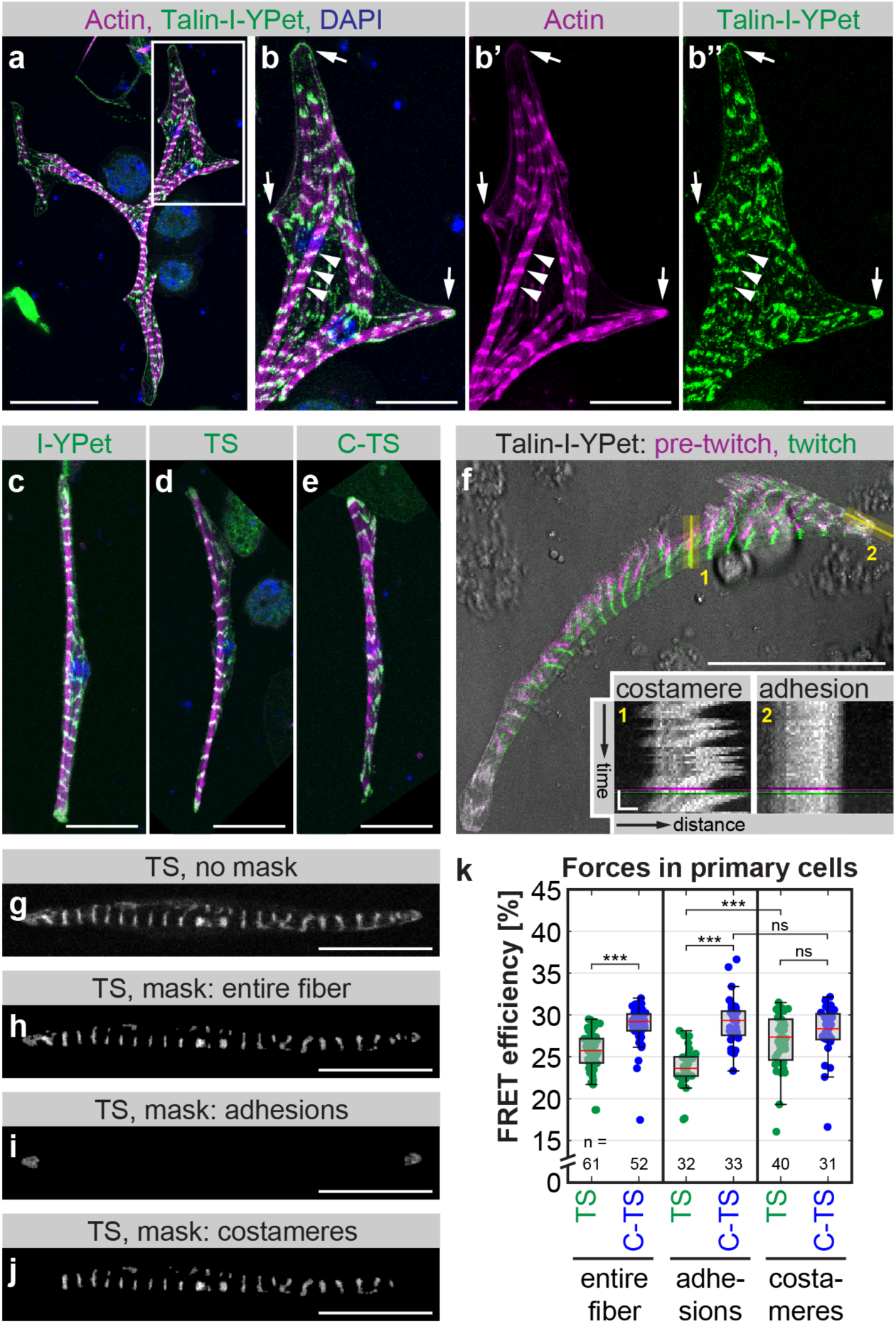
Talin tension sensor reveals forces in primary muscle fibers. **a-b**, Primary myoblasts isolated from *talin-I-YPet* embryos were differentiated and stained with phalloidin and DAPI on day 6. White box in **a** indicates zoom-in area in **b**. In differentiated muscle fibers Talin-I-YPet localizes to adhesions at fiber tips (arrows) and to costameres along myofibrils (arrowheads). **c-e**, Primary muscle fibers differentiated from *talin-I-YPet*, *talin-TS*, or *talin-C-TS* embryos stained with phalloidin (magenta) and DAPI (blue) show similar morphologies and Talin localisation (green). **f**, Transmission light image (grey) of a twitching primary muscle cell overlaid with Talin-I-YPet signal pre-twitch (magenta) and during the twitch (green), and kymographs of the regions indicated in yellow. Note that costameres move with contractions while adhesions are fixed to the substrate. **g-j**, Masking of cells for force analysis. From the original image (**g**) masks from the entire fiber (**h**), from adhesions at fiber tips (**i**) or from costameres (**j**) were created. **k**, Talin forces measured by FLIM-FRET. A decrease in FRET efficiency of Talin-TS (TS) compared to the C-terminal zero-force control (C-TS) indicates force. Note that Talin in adhesions experiences a significant amount of force while Talin in costameres does not (Kolmogorov-Smirnov test, *** p<0.001, ns=not significant p>0.05; n=number of fibers). Scale bars are 50 μm in **a** and **f** and 20 μm in **b-e** and **g-j**. Scale bars in kymographs in **f** are 10 s and 2 μm.

For establishing force measurements using these primary fiber cultures, we performed fluorescence lifetime imaging microscopy (FLIM) to determine the FRET efficiency of the Talin tension sensor containing the HP sensor module (TS) compared to the zero-force control (C-TS). We created distinct masks for Talin FRET signals either in the entire fiber, or only in cell-substrate adhesions at the fiber tips, or in costameres along myofibrils (Fig. 2g-j). Consistent with earlier Talin force measurements, we observed a reduction in FRET efficiency of TS compared to the control C-TS within the entire fiber, indicating that Talin indeed experiences mechanical forces in these adherent, primary muscle fibers. As expected, we find higher average forces across Talin at muscle-substrate adhesions compared to the rest of the cell. In costameres, which are not fixed to the plastic substrate, the FRET efficiency of TS is indistinguishable from the control, indicating that forces across Talin at costameres are lower and do not exceed 6-8 pN. Together, these data demonstrate that the *Drosophila* Talin tension sensor reports similar Talin forces at adhesions of cultured muscle fibers as were previously described for Talin in focal adhesions of mammalian fibroblasts [11,12,16].

### Forces across *Drosophila* Talin *in vivo*

To quantify forces across Talin *in vivo*, we chose the developing muscle-tendon attachments of the flight muscles as a model system. At 20 hours after puparium formation (h APF), the developing myotubes have initiated contact with the tendon epithelium and immature muscle attachment sites are formed (Fig. 3a). At 24 h APF, the attachment sites have started to mature while the myotubes compact and long cellular extensions from the tendon epithelium are formed. During this process, increasing mechanical tension is build up in the tissue [23]. At 30 h APF, the myotubes reach their maximally compacted stage and initiate myofibrillogenesis, before the muscles elongate and grow to fill the entire thorax by the end of the pupal stage [28]. Since muscle attachment critically depends on integrins and Talin function to resist tissue tension [23,30], it is an ideally suited developmental system to investigate molecular forces across Talin *in vivo*.

**Fig. 3.**
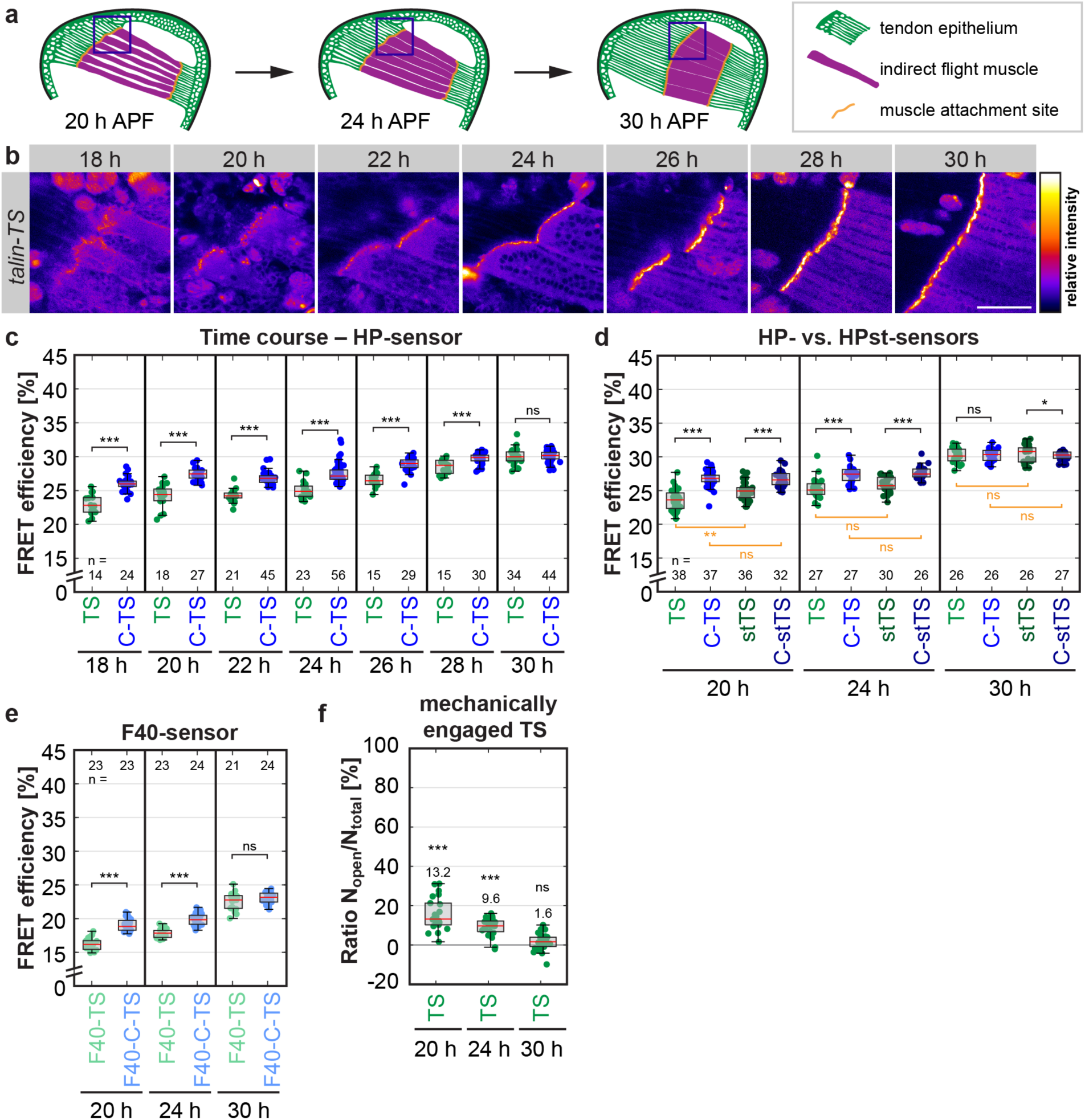
A small proportion of Talin molecules at muscle attachment sites *in vivo* are mechanically engaged. **a**, Schemes of indirect flight muscle development in the pupal thorax at 20, 24 and 30 h APF. Blue boxes indicate areas imaged for force measurements (see **b**). **b**, Images showing Talin tension sensor (TS) localization to maturing muscle attachment sites. Scale bar is 50 μm. **c**, Talin forces measured by FLIM-FRET in a time course using the HP-sensor module (6-8 pN). A decrease in FRET efficiency of Talin-TS (TS) compared to the C-terminal zero-force control (C-TS) indicates force. Note that the average force per molecule is highest in the beginning of the time course. **d**, Comparison of TS (6-8 pN) and stTS (9-11 pN) to the C-terminal zero-force controls, C-TS and C-stTS. Note that both sensors indicate forces across Talin at 20 h and 24 h APF (significance indicated in black). Direct comparisons between TS and stTS or the controls are indicated in orange. Note the increase in FRET of stTS compared to TS at 20 h APF. **e**, Talin force measurements using the F40-sensor module (1-6 pN). **f**, Proportion of mechanically engaged TS determined as the ratio of open (N_open_) vs. total sensor (N_total_) using biexponential fitting. Significance is indicated in comparison to zero-force control level (set to zero). The raw data are the same as in **c**. (Kolmogorov-Smirnov test, *** p<0.001, ** p<0.01, * p<0.05, ns=not significant p>0.05; n=number of pupae).

We measured Talin forces between 18 and 30 h APF in living pupae at the anterior muscle attachment sites of the dorsal-longitudinal flight muscles using the HP-sensor module (Fig. 3b and workflow in Extended Data Fig. 2). For calculating the FRET efficiency, we determined the donor fluorescence lifetime in flies expressing YPet alone at the internal position of Talin (I-YPet) (Extended Data Fig. 3a). In addition, we excluded the possibility that FRET between neighbouring molecules (intermolecular FRET) affects our measurements throughout the entire time course (Extended Data Fig. 3b) and confirmed that our lifetime measurements are independent of signal intensity (Extended Data Fig. 3c). When we compared FRET efficiencies in *talin-TS* and *talin-C-TS* animals, we detected a significant drop in FRET efficiency for Talin-TS at 18-28 h APF. However, the FRET efficiency reduction at muscle attachment sites was significantly smaller compared to the *in vitro* measurements of cultured muscle fibers (Fig. 2k) or of cultured mammalian fibroblasts [11]. At 30 h APF, no difference in FRET efficiencies was detected, suggesting that there is little or no tension across Talin at this time point. Together, these data suggest either that forces per Talin molecule are largely below 6-8 pN or that only a small percentage of Talin molecules at muscle attachments experience forces above 6 pN at 18-28 h APF. Contrary to our expectation, the average force across Talin decreases during muscle compaction when tissue tension is known to build up and myofibrils are assembled.

To substantiate these findings, we compared flies carrying the HP-based Talin sensor (6-8 pN) to those with the stable variant HPst (9-11 pN), which only differs in two point mutations. We found similar and highly reproducible differences in FRET efficiency (Fig. 3d, Extended Data Fig. 3d) indicating that at 20-24 h APF, some Talin molecules experience forces of even ≥10 pN at muscle attachment sites. Importantly, comparison of TS to its stable variant (stTS) revealed a significant difference in FRET efficiency at 20 h APF while the respective zero-force controls were indistinguishable (Fig. 3d). This demonstrates that a proportion of the mechanically engaged Talin molecules experience a range of forces between 7 and 10 pN at muscle-tendon attachments *in vivo*, further emphasizing that the observed differences are force-specific.

To test whether the remaining Talin molecules experience forces that are too low to be detected by the HP or HPst sensor modules, we generated flies with the F40 sensor module, which is sensitive to forces of 1-6 pN [13]. Again, we quantified a decrease in FRET efficiency relative to the control at 20 h and 24 h APF but FRET efficiency differences remained small and no change was observed at 30 h APF (Fig. 3e). Thus, a large proportion of the Talin molecules at muscle attachment sites are not exposed to significant mechanical forces during development.

To quantify the proportion of mechanically engaged Talin molecules at 20 h and 24 h APF, we applied biexponential fitting to our FLIM data and calculated the ratio of open *vs*. closed sensor (Fig. 3f, see methods for details). This analysis revealed that only 13.2% and 9.6% of all Talin molecules are mechanically engaged at 20 h and 24 h APF, which contrasts *in vitro* measurements of focal adhesions that are characterized by a Talin engagement ratio of about 70% [12].

### Talin concentration at developing muscle attachments

Since Talin is thought to play an important mechanical role during tissue formation, we wanted to test whether such a small proportion of mechanically engaged Talin molecules *in vivo* could still contribute a significant amount of tissue-level tension. We therefore quantified the absolute amount of Talin molecules present at muscle attachment sites by combining *in vivo* fluorescence correlation spectroscopy (FCS) with quantitative confocal imaging (see workflow in Extended Data Fig. 4a-d). From FCS measurements in the muscle interior we calculated the counts per particle (CPP) value, i.e. the molecular brightness of a single Talin-I-YPet particle in each pupa. Since such a particle may correspond to a Talin monomer or dimer, we compared the Talin-I-YPet brightness to the brightness of free monomeric YPet expressed in flight muscles and found no significant difference (Fig. 4a). We conclude that Talin is mostly monomeric in the muscle interior.

**Fig. 4.**
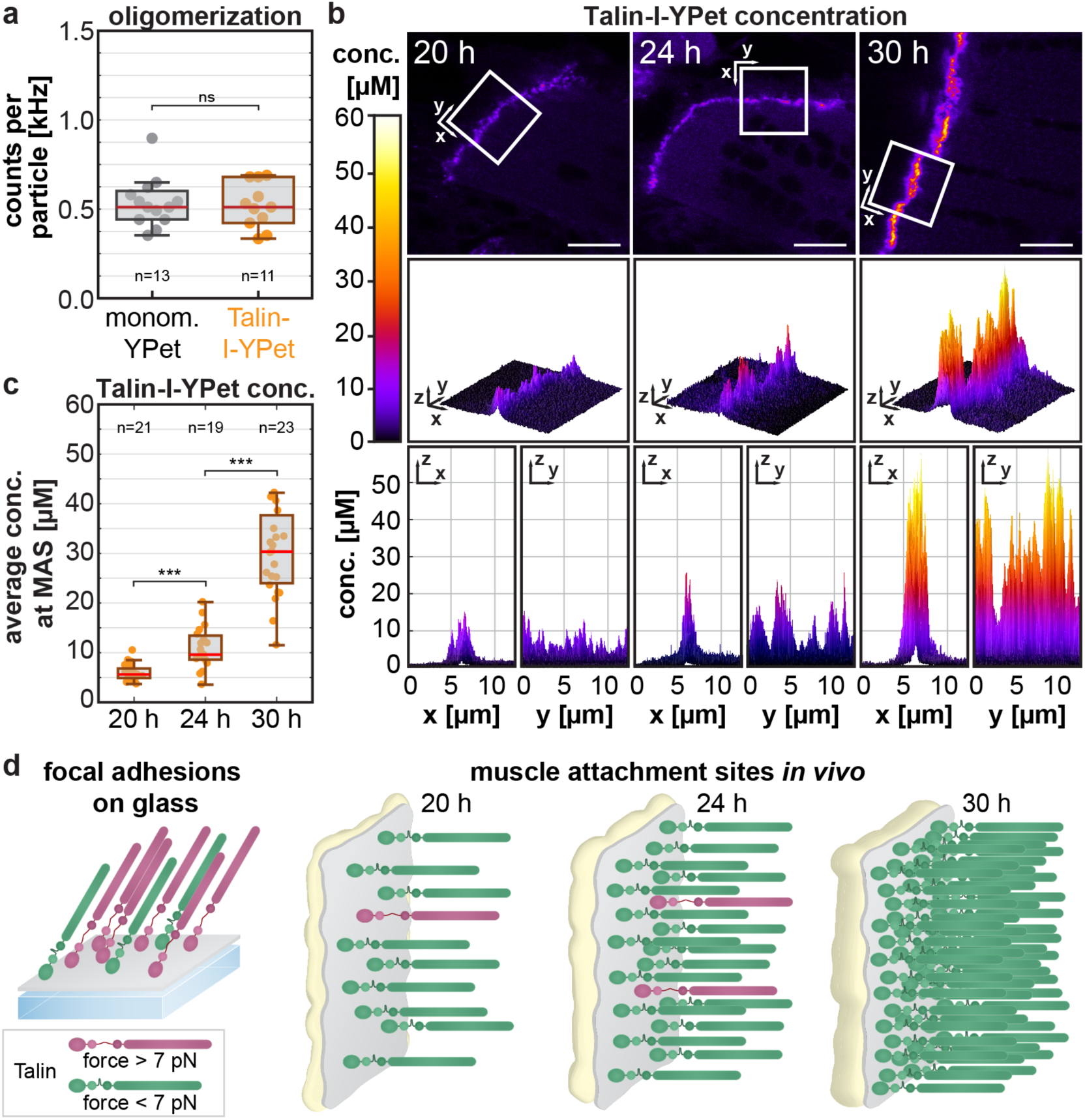
Talin concentration at muscle attachment sites increases five-fold during attachment maturation. **a**, Degree of Talin oligomerization measured by *in vivo* fluorescence correlation spectroscopy (FCS) in the muscle interior. Brightness (in counts per particle) of monomeric free YPet compared to Talin-I-YPet particles. Note that Talin-I-YPet particles are as bright as monomeric YPet, thus Talin-I-YPet is also monomeric (Kolmogorov-Smirnov test, ns=not significant p=0.976, n=number of pupae). **b**, Absolute Talin-I-YPet concentration (conc.) measured by FCS in combination with quantitative confocal imaging. Representative calibrated concentration images are shown for 20, 24 and 30 h APF. The boxes mark the area shown in the graphs below from different perspectives as indicated. Scale bars are 10 μm. **c**, Quantification of the average Talin-I-YPet concentration at the muscle attachment sites (MAS) per image. Note that the concentration increases about 2-fold from 20 to 24 h APF and 5-fold to 30 h. (Kolmogorov-Smirnov test, *** p<0.001, n=number of pupae) **d**, Model of mechanical Talin engagement. In focal adhesions, 70% of the Talin molecules are under force[12] while at muscle attachment sites *in vivo* less than 15% are mechanically engaged at any given time. As more Talin is recruited during muscle attachment maturation, the proportion of mechanically-engaged Talin molecules decreases even further.

Next, we calculated the Talin concentration a muscle attachment sites by calibrating confocal images using the molecular brightness (CPP) information from the FCS measurements. Using a dilution series of Atto488, we ascertained that the fluorescence intensity increases linearly with the concentration over multiple orders of magnitude in our measurements (Extended Data Fig. 4e). The resulting images with pixel-by-pixel Talin concentration values (Fig. 4b) indicate an average concentration at the muscle attachment of 5.9 μM (20 h), 10.9 μM (24 h) and 30.9 μM (30 h) (Fig. 4c). Thus, the local concentration of Talin molecules increases approximately two-fold from 20 h to 24 h and five-fold to 30 h, indicating that Talin may contribute to the overall increase in tissue stress by its strong recruitment to maturing muscle attachment sites.

To confirm this hypothesis, we estimated the density of Talin molecules on the membrane by dividing the number of Talin molecules per pixel by the estimated membrane area in the confocal volume (Fig. 4d, see Methods for details). This resulted in about 400, 700 and 2300 Talin molecules per μM^2^ at 20 h, 24 h and 30 h APF, respectively, which corresponds to 20 nm × 20 nm space per molecule at 30 h APF. This space can easily accommodate the size of a Talin head domain (about 4 nm × 10 nm) [34], and the estimated density is comparable to previous studies of integrins in focal adhesions [35].

By combining our force quantifications with the estimated Talin density at muscle attachment sites, we calculated the Talin-mediated tissue stress to be in the order of 0.40-0.5 kPa at 20-24 h APF (see methods for details). These values are remarkably close to a previously published stress estimate of 0.16 kPa determined by traction force microscopy in focal adhesions of cultured cells [36]. Thus, Talin does contribute a significant amount of tissue stress despite the small proportion of mechanically engaged molecules (Fig. 4d).

## Discussion

Our findings highlight the importance of investigating tissues in their natural mechanical environment *in vivo*. While the forces per Talin molecule and the tissue stress *in vivo* are in the same order of magnitude as in previous *in vitro* studies of focal adhesions [11,12,36], a surprisingly small proportion of Talin molecules (<15%) experience detectable forces during muscle development *in vivo*. An obvious question arising from our finding is: What are the other Talin molecules doing at muscle attachment sites, for which we cannot detect significant mechanical forces? Likely, the pool of mechanically engaged Talin molecules exchanges dynamically with the other Talin molecules present at the muscle attachment site. Such a dynamic system can allow the attachment to rapidly adjust to changes in tissue forces preventing rupture of the muscle-tendon attachment upon a sudden increase in tissue force.

It has been shown previously in focal adhesions of cultured cells, that the length of Talin can fluctuate dynamically on the time scale of seconds, with Talin being transiently extended from 50 nm up to 350 nm [37]. This can be explained with reversible folding and unfolding of some of the 13 helical bundles in the Talin rod upon actomyosin-dependent stretching of Talin. The unfolding of the helical bundles makes binding sites accessible leading to recruitment of vinculin [38]. The dynamic stretching of Talin could allow it to act as a ‘shock-absorber’ [25], consistent with our finding that only some molecules are under force at the same time, while additional molecules could be present to make the attachment robust against higher forces.

The forces required to unfold the Talin rod domains to enable vinculin binding are well within the range of the force sensors used here. The rod domain R3 unfolds at about 5 pN [39] and the remaining rod domains unfold when forces larger than 8 pN are applied [40]. Hence, our force sensors detect biologically significant forces that change the Talin structure making vinculin binding sites accessible and allowing a mechanotransduction response.

Previous estimates of forces transmitted by integrins based on studies of focal adhesions *in vitro* cover a wide range of forces. Studies using extracellular sensors with synthetic integrin ligands (that report forces based on double-stranded DNA rupture) suggest that integrins can experience very high forces in cells plated on glass (more than 53 pN) [41,42]. However, other data generated with FRET-based extracellular sensors suggest that about 70% of the integrins in focal adhesions experience low forces (less than 3 pN) [43]. While these *in vitro* systems have the advantage that they are accessible for precise manipulations, the artificial mechanical environment may have a strong impact on the amount of force experienced by the individual proteins and the number of molecules that are mechanically engaged. Our study provides, to our knowledge, the first insights into molecular forces acting on integrin-mediated attachments *in vivo*. Here we focus on developing muscle attachments in pupae, notably the built reagents should enable future force measurements in all integrin-based processes in *Drosophila*. Based on our finding that only a small proportion of Talin molecules (<15%) are experiencing forces higher than 6-8 pN at muscle attachments, we hypothesize that tissues prevent mechanical failure during development *in vivo* with the following mechanism: a large pool of molecules dynamically share the mechanical load, such that a sudden increase in tissue tension can be rapidly buffered by mechanically engaging additional molecules already present at the attachment site. Mechanical failure of integrin-mediated attachments *in vivo* needs to be avoided at all cost, particularly in muscle fibers or cardiomyocytes, to prevent fatal consequences for the animal. Hence, creating a buffer to withstand peak forces can be an important concept for the survival of animals.

## Methods

### Fly strains

All fly work was performed at 27°C to be consistent with previously published work, unless otherwise stated. For details on the genome engineering strategy resulting in Talin tension sensor and control stocks generated in this study (*talin-F40-TS*, *talin-C-F40-TS*, *talin-TS*, *talin-C-TS*, *talin-stTS*, *talin-C-stTS*, *talin-I-YPet*, *talin-C-YPet*, *talin-I-mCh*, and *talin-C-mCh*) see below. Other stocks used were *Mef2*-GAL4 [44] and *UAS*-mCherry-Gma [45].

### Generation of tension sensor and control stocks

Tension sensor and control stocks were generated by combining CRISPR/Cas9-mediated genome engineering with recombinase-mediated cassette exchange (RCME) as described previously [27]. See Extended Data Fig. 1 for a detailed depiction of the 2-step strategy. For step 1, single guide (sg)RNAs were designed with the help of an online tool maintained by the Feng Zhang lab (http://crispr.mit.edu/) [46] and transcribed *in vitro*. After testing sgRNA cutting efficiency in Cas9-expressing S2-cells [47], two sgRNAs (70 ng/μL) were injected into *Act5C-Cas9*, *Lig4^169^* embryos together with the dsRed donor vector (500 ng/μL) containing a dsRed eye marker cassette flanked by attP sites and homology arms. Successful homologous recombination events were identified by screening for red fluorescent eyes and verified by PCR and sequencing. “Ends-in” events were excluded. We call the resulting fly lines *talin-I-dsRed and talin-C-dsRed*. For step 2, vasa-ϕC31 plasmid (200 ng/μL) was injected together with attB-donor vector (150 ng/μL). Successful exchange events were identified by screening for the absence of dsRed and correct orientation of the cassette was verified by PCR.

### Adult hemithorax staining

Adult hemithoraxes were dissected and stained in a similar way as previously described [48]. Specifically, the wings and abdomen were cut off the thorax of adult males (1 day old) with fine scissors and the thoraxes were fixed for 15 min in 4% PFA in relaxing solution (20 mM sodium phosphate buffer, pH 7.0; 5 mM MgCl_2_; 5 mM ATP; 5 mM EGTA; 0.3% Trition-X-100). After washing once with PBST (PBS with 0.3% Triton-X-100), the thoraxes were placed on double-sided tape and the legs were cut off. Next, the thoraxes were cut sagittally with a microtome blade (dorsal to ventral). The thorax halves were placed in PBST, washed once and blocked in normal goat serum (1:30) for 30 min at room temperature (RT) on a shaker. Primary antibodies (anti-Talin antibody: 1:500, 1:1 mixture of E16B and A22A, DSHB) were incubated overnight at 4°C on a shaker. Hemithoraxes were then washed 3x 10 min in PBST at RT and stained with secondary antibody (Alexa488 goat anti-mouse IgG, 1:500, Molecular Probes) and phalloidin (Rhodamine- or Alexa647-conjugate, 1:500 or 1:200 respectively, Molecular Probes) in PBST for 2 hours at RT in the dark. After washing 3x with PBST for 5 min, hemithoraxes were mounted in Vectashield containing DAPI with two spacer coverslips on each side. YPet signal after fixation was bright enough for imaging without further amplification.

### Dissection of pupae

32 h APF pupae were freed from the pupal case and dissected in PBS in a silicone dish using insect pins [48]. The head and the sides were cut using fine scissors to remove the ventral half of the pupa. Next, the thorax was cut sagittally and the thorax halves were cut off the abdomen and placed in fixing solution (4% PFA in PBST) for 15 min. The thorax halves were then stained with phalloidin and DAPI like the adult hemithoraxes but without shaking and mounted using one spacer coverslip.

### Imaging of stainings

Samples were imaged on a Zeiss LSM 780 scanning confocal microscope with Plan Apochromat objectives (10x air, NA 0.45 for overview images and 40x oil, NA 1.4 for detail images). For thick samples, a z-stack was acquired and maximum-projected using ImageJ.

### Sarcomere length quantification

Sarcomere length was quantified as previously described using the ImageJ plug-in MyofibrilJ (https://imagej.net/MyofibrilJ) [28]. Briefly, an area with straight, horizontal myofibrils is analysed by Fourier transformation to find the periodicity of the sarcomeres. One area was analysed for each hemithorax stained with phalloidin and imaged at 40x and zoom 4.

### Western blotting

Western blotting was performed according to standard procedures. Specifically, 15 flies each were homogenized in 100 μL 6x SDS loading buffer (250 mM Tris pH 6.8, 30% glycerol, 1% SDS, 500 mM DTT) and heated to 95°C for 5 min. 200 μL of water were added and the equivalent of 0.5 (10 μL) and 1 fly (20 μL) were loaded onto a NuPAGE Novex 3-8% Tris-Acetate Gel. The transfer to the membrane was carried out overnight. The membrane was blocked (5% blotting grade blocker, BioRad) and then incubated overnight with a 1:1 mixture of anti-Talin antibodies E16B and A22A (1:1000 in block). For detection, HRP anti-mouse antibody and Immobilon Western Chemiluminescent HRP Substrate (Millipore) were used.

### Flight assays

Male flies (1-3 days old, aged at 25 °C) were thrown into a 1 m × 8 cm plexiglass cylinder with 5 marked sections [49]. Flightless flies fall to the bottom of the tube immediately while strong fliers land in the top two sections and weak fliers in the 3^rd^ and 4^th^ section. Flight assays were performed in triplicates with 10-20 males each and repeated twice.

### Live imaging of embryos and larvae

Embryos from the cross *yw;; talin-I-YPet* to *w; Mef2-GAL4; UAS-mCherry-Gma* were collected on apple juice agar plates for 24 hours and dechorionated in 50% bleach (0.024% hypochlorite) for 3 min. Living embryos were mounted in 50% glycerol before imaging. L3 larvae from the same cross were immobilised by immersing them in 60°C water for about 1 s [29] and mounted using a plexiglass slide with a groove and one spacer coverslip on each side in 50% glycerol. 5×1 tile scan z-stacks were acquired using a 10x objective to image the entire larva.

### Isolation and differentiation of primary muscle fibers

Primary cells were isolated from *Drosophila* embryos and differentiated as previously described [31,32] with the following modifications: Embryos (5-7 hours old, aged at 25°C) were collected from smaller cages on only one 9 cm molasses plate per genotype. Embryos were homogenized with a Dounce homogeniser using a loose fit pestle in 4 mL Schneider’s *Drosophila* medium (Gibco 21720-024, lot 1668085) and after several washing steps (using 2 mL medium) re-suspended to a concentration of 3×10^6^ cells/mL. Finally, cells were plated in 8-well ibidi dishes (1 cm^2^ plastic bottom for microscopy with ibiTreat surface) coated with vitronectin (optional) at a density of 3-9×10^5^ cells/cm^2^ and differentiated for 5-7 days at 25°C in a humid chamber.

### Fixation, staining and imaging of primary muscle fibers

Primary muscle fibers were fixed on day 6 after isolation with 4% PFA in PBS for 10 min at RT on a shaker. Phalloidin-staining (Alexa647-conjugate, Molecular Probes) was performed overnight in the dark at 4°C. Fixed cells were imaged in PBS on a Zeiss LSM 780 with a 40x oil objective (Plan Apochromat, NA 1.4). Live imaging of twitching primary cells was performed on a Leica SP5 confocal with a 63x water objective (HCX PL APO 63x/1.2 W CORR λ_BL_), acquiring the transmission light channel and the YPet channel simultaneously.

### Sample preparation for live imaging of pupae

White pre-pupae were collected and aged at 27°C to the desired time point. Before imaging, a window was cut into the pupal case above the thorax and the pupae were mounted on a custom-made slide with a groove as previously described [50].

### Fluorescence lifetime imaging microscopy (FLIM)

Primary muscle fibers and pupae were imaged live on a Leica SP5 microscope equipped with a pulsed white light laser (NKT Photonics, 80 MHz), a time-correlated single photon counting (TCSPC)-FLIM detector (FLIM X16, LaVision BioTec) and a 545/30 nm emission filter (Chroma). Primary muscle fibers were imaged with a 63x water objective (HCX PL APO 63x/1.2 W CORR λ_BL_) and pupae were imaged with a 40x water objective (HC PL APO 40x/1.1 W CORR CS2). Photon arrival times were detected with a resolution of 0.08 ns in a 12.5 ns time window between laser pulses.

### FLIM-FRET data analysis

The FLIM data were analysed using a custom-written MATLAB program [11,12]. First, an intensity image was created to manually draw a region of interest (ROI) around the target structure (adhesions/costameres in primary cells or muscle attachment sites in pupae, also see Extended Data Fig. 2). To create a binary mask of the target structure, Multi-Otsu thresholding with three classes was applied to the signal in the ROI blurred with a median filter (3×3 pixels) and holes in the mask containing the brightest class were filled. Photon arrival times of all photons inside the mask were plotted in a histogram and the tail of the curve was fitted with a monoexponential decay yielding the fluorescence lifetime *τ*. Fits with more than 5% relative error in lifetime determination were excluded from further analysis. For dimmer samples (primary fiber cultures and intermolecular FRET pupae), we used a 10% relative error cut-off. The FRET efficiency *E* was calculated according to the following formula with *τ_DA_* being the lifetime of the donor in presence of the acceptor and *τ_D_* the lifetime of the donor alone:

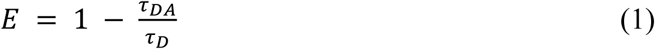

For all measurements, *τ_D_* was determined using Talin-I-YPet. Experiments were repeated 2-5 times on different experiment days with 10-15 pupae/cells imaged per genotype and day.

### Calculation of the proportion of mechanically engaged Talin

We determined the number of mechanically engaged (=open) tension sensor *N_open_* relative to the total number of molecules *N_total_* at the muscle attachment site using biexponential fitting similarly as previously described [12]. Briefly, we assumed that the fluorescence decay from a tension sensor FLIM measurement can be described by two lifetimes: The lifetime of the open sensor *τ_no_FRET* and the lifetime of the closed sensor undergoing FRET *τ_FRET_*. The lifetime of the open sensor *τ_no_FRET* approximately corresponds to the lifetime of the donor alone, because of the large contour length increase upon opening of the sensor. Thus, we determined the lifetime *τ_no_FRET* by using a monoexponential fit on Talin-I-YPet data as described above. The lifetime *τ_FRET_* was determined from zero-force control (Talin-C-TS) data. Since the Talin-C-TS sample contains fully fluorescent sensor (*τ_fret_*) and sensor with non-fluorescent mCherry acceptor (*τ_noFRET_*), we used a biexponential fit with fixed *τ_no_FRET* to determine *τ_FRET_*. The two lifetimes *τ_no_FRET* and *τ_FRET_* were then fixed and used to fit Talin-TS and Talin-C-TS data biexponentially, thereby determining the relative contributions of photons from molecules with these two lifetimes. From this, the relative number of molecules with *τ_no_FRET* and *τ_FRET_* was calculated, taking into account that FRET reduces the number of photons detected in the donor channel. Finally, the ratio *N_open_*/*N_total_* was determined by normalizing the Talin-TS values to the respective Talin-C-TS values.

### Fluorescence correlation spectroscopy (FCS)

Living *talin-I-YPet* pupae were analysed at 20 h, 24 h and 30 h APF by a combination of confocal microscopy (LSM 780, Zeiss) and FCS using a 40x water immersion objective (C-Apochromat 40x/1.20 W Korr UV-VIS-IR) and the built-in GaAsP detector in single photon counting mode. Prior to experiment, the correction collar and pinhole position were adjusted with fluorescent Rhodamine 6G in aqueous solution (30 nM in Tris pH 8) using the same type of cover glass (Marienfeld, High Precision, 18×18 mm, 170±5 μM thickness) as for mounting the pupae [50]. To calibrate the detection volume (excitation 514 nm laser light), we measured FCS (120 s recordings) at three different positions 20 μM above the cover glass surface. Autocorrelation curves were analysed with our open-source software *PyCorrFit* [51] (Version 1.0.1, available online at http://pycorrfit.craban.de/). For fitting Rhodamine 6G data we used a model accounting for triplet transitions and three-dimensional diffusion (denoted “T-3D” in *PyCorrFit*). The detection volume *V_eff_* was calculated based on the measured diffusion time (*τ_diff_*) and the published diffusion coefficient D = 414 μM^2^/s [52]:

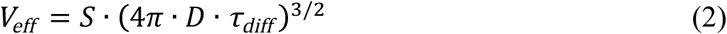

For all measurements, the axis ratio of the detection volume S = 5 was consistently fixed [53]. In living pupae, fluorescent proteins (YPet or Talin-I-YPet) were measured by FCS using a park and probe procedure [54]: In images, three positions in the muscle interior next to the muscle attachment site were manually selected for FCS (10x 40s recordings). For fitting of Talin-I-YPet autocorrelation curves (time bins > 1 μs), a two-component three-dimensional diffusion model with two non-fluorescent dark states (denoted “T+T+3D+3D” in *PyCorrFit)* was applied. Transient dark states were assigned either to triplet transitions (τ_trip1_, T_1_) in the time range of 1-20 μs and photochemical flickering (τ_trip2_, T_2_) in the time range of about 200-600 μs [55]. The first diffusion time was assigned to protein diffusion in the muscle interior whereas the second diffusion term was merely a descriptive term accounting for slow long tail behaviour that cannot be avoided in a crowed intracellular environment [54]. Autocorrelation curves derived from visibly unstable intensity traces were excluded from further analysis. Due to the high endogenous expression levels, the contribution of non-correlated background was negligible. Thus, the molecular brightness, i.e. the counts per particle (CPP) value of Talin-I-YPet was determined by dividing the average intensity *I* (brackets indicate the average) by the number of molecules in the focal volume *N*, which is dependent on the autocorrelation amplitude G(0) (of the autocorrelation function G(τ)) and the dark fractions T_1_ and T_2_ from the fit:

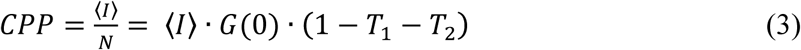

Since freely diffusing YPet diffuses faster than Talin-I-YPet, the signal fluctuations related to flickering and diffusion cannot be distinguished in YPet measurements. Therefore, the autocorrelation curves of free YPet were fitted by a simplified model function accounting only for transient triplet states and two diffusive terms, of which the first combines contributions of both protein diffusion and flickering (denoted “T-3D-3D” in *PyCorrFit*). To estimate true particle numbers, we corrected for triplet transitions and flickering globally by using the average fractions T_1_ and T_2_ from corresponding Talin-I-YPet measurements performed with the same excitation power density:

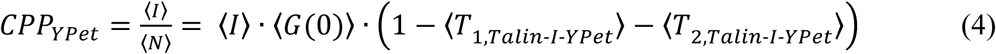

The diffusion constant of freely expressed YPet was in good agreement to other fluorescent proteins in the cytoplasm of living cells, suggesting the point spread function positioned in the muscle cell is still diffraction limited. This finding justifies the external calibration of the detection volume by Rhodamine 6G.

### Calibration of confocal images

For quantification of the Talin-I-YPet concentration at muscle-tendon attachment sites, the developing flight muscles were imaged in photon counting mode (512×512 px, pixel dwell time *PT*=50 μs). Saturation of the detector was carefully avoided (*I*(x,y) < 2 MHz). The counts in each pixel of an image were calibrated by the molecular brightness (CPP) value determined for Talin-I-YPet in the interior of the same muscle fiber by FCS[54]. Due to the monomeric state of Talin-I-YPet, intensity values stored in each pixel *I*(x,y) could be directly transformed into numbers of Talin molecules:

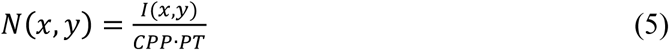

Using the Avogadro constant and the detection volume (*V_eff_*) as determined by Rhodamine 6G measurements, we then calculated concentration maps:

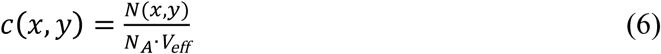

Finally, the muscle attachment sites were isolated in the Talin-I-YPet concentration maps by creating a mask with the same thresholding algorithm as used for FLIM-FRET. The concentration values were averaged across pixels within the mask resulting in a mean concentration value per pupa.

### Estimation of Talin density and tissue stress

To estimate Talin density on the membrane from pixel-by-pixel concentration values, we divided the average number of molecules in the focal volume at the muscle attachment sites by the membrane area in the focal volume. The focal volume was determined by Rhodamine 6G FCS measurements as described above. For the shape of the focal volume we assumed an ellipsoid with the long axis (z) being 5-times the short axis (x=y). Hence, for a focus volume of 0.32 fL, the membrane area in the z-y-plane is 0.63 μM^2^. Taking into account that there are two membranes (one from the tendon and one from the muscle) and that the membrane is not flat (ruffles approximately increase the area 2-fold as determined from EM-images [56]) the total membrane area in the focal volume is about 2.5 μM^2^.

To estimate Talin-mediated tissue stress, we calculated *force threshold of sensor* × *Talin density* × *proportion of mechanically engaged Talin* = 7 pN × 400 molecules/μm^2^ × 13.2% = 0. 37 kPa for 20 h APF and 7 pN × 700 molecules/μm^2^ × 9.6% = 0.47 kPa for 24 h. Note, that these values are lower estimates since individual molecules might experience forces higher than 7 pN.

### Statistics

Box plots display the median as a red line and the box denotes the interquartile range. Whiskers extend to 1.5 times the interquartile range from the median and are shortened to the adjacent data point (Tukey). In addition, all data points are shown as dots. Tests used for statistical evaluation are indicated in the figure legends.

### Code availability

FLIM-FRET data was analysed using custom-written MATLAB code as published previously [11,12]. The code is available upon request.

## Acknowledgements

This work was supported by the EMBO Young Investigator Program (F.S.), the European Research Council under the European Union’s Seventh Framework Programme (FP/2007-2013)/ERC Grant 310939 (F.S.), the Max Planck Society (S.B.L., T.W., A.-L.C., CG, F.S.), the Centre National de la Recherche Scientifique (CNRS) (F.S.), the excellence initiative Aix-Marseille University AMIDEX (ANR-11-IDEX-0001-02, F.S.), the LabEX-INFORM (ANR-11-LABX-0054, F.S.), the ANR-ACHN (F.S.), the Human Frontiers Science Program (HFSP, F.S.), the Bettencourt Foundation (F.S.), the Boehringer Ingelheim Fonds (S.B.L.), the German Research Council (DFG) priority program SPP1782 (C.G.) and a Human Frontier Science Program Grant (RGP0024, C.G).

The authors are indebted to Carleen Kluger (initial software development for FLIM-FRET data analysis), Paul Müller (PyCorrFit software development), Bettina Stender, Nicole Plewka, Christophe Pitaval and Céline Guichard (fly embryo injections), Xu Zhang (two-step CRISPR/RMCE protocol), Petra Schwille (access to FCS-equipment) and Reinhard Fässler (continuous support).

## Author contributions

S.B.L. performed all the experiments, with support from T.W. for the FCS experiments. S.B.L. analysed all the data with help from A.-L.C. and generated the figures. A.-L.C. refined FLIM analysis software. F.S. conceived and supervised the project with essential input from C.G. throughout the project. S.B.L. and F.S. wrote the manuscript with input from all authors.

## Data availability statement

The authors declare that the relevant data supporting the findings of this study are included within the paper. Additional data are available upon request to the corresponding authors.

## Correspondence & Materials

Correspondence and requests for materials should be addressed to F.S, C.G or S.B.L.

---

**Extended Data Fig. 1.**
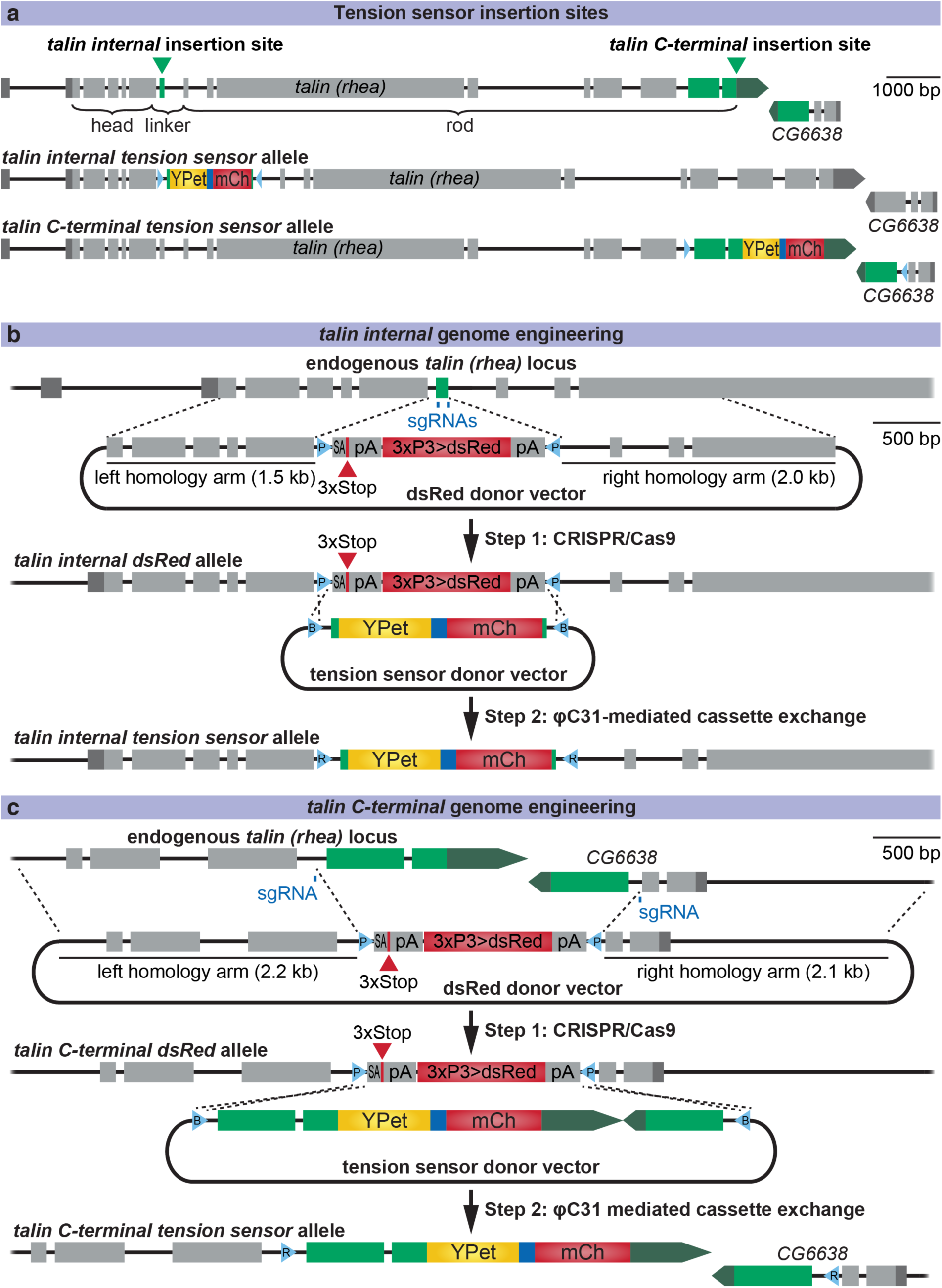
Talin tension sensor genome engineering. **a, Top:** Gene model of *talin* (*rhea*, *isoform RF*) with the insertion sites (green) in the linker region between Talin head and rod (internal), and at the C-terminus. The gene *CG6638* immediately follows *talin* and therefore was included, too. **Middle:** Tension sensor allele with the sensor module inserted into the target exon in the linker region of Talin. attR sites left in the surrounding introns are shown in light blue. **Bottom:** C-terminal control sensor allele with the sensor module inserted at the C-terminus of Talin. Gene models are drawn to scale. **(b)** Scheme showing how tension sensor alleles were generated. **Step 1:** The target exon in the linker region (green) was replaced by a splice acceptor (SA)-3xStop-SV40 terminator (pA)-3xP3>dsRed-pA cassette flanked by attP sites (P) using the CRISPR/Cas9 system. Specifically, a dsRed donor vector containing 1.5 to 2.0 kb homology arms was injected into *Act5C*-Cas9 expressing embryos (also carrying a *lig4^169^* mutation to favour homology directed repair over non-homologous end-joining [27]) together with two *in vitro*-transcribed single guide (sg)RNAs (target sites in blue). Successful targeting was identified by screening for fluorescent red eyes. **Step 2:** ϕC31-mediated cassette exchange was performed to replace the dsRed cassette by the original target exon including a tension sensor module consisting of YPet, a flexible calibrated, mechano-sensitive linker peptide (dark blue) and mCherry (mCh). To this end, a tension sensor donor vector including flanking attB sites (B) was injected together with vasa-ϕC31 plasmid. Thereby, the tension sensor was inserted seamlessly into the gene (after Talin amino acid 456) except for two attR sites (R) in the flanking introns. Successful exchange events were identified by screening for the absence of fluorescent red eyes [27]. Control fly lines with one fluorophore and fly lines with different tension sensor modules were generated by repeating step 2 with different donor vectors. **(C)** Scheme showing how C-terminal zero-force sensor alleles were generated using the same strategy. However, at the C-terminus three exons (green) were replaced by the dsRed cassette in the first step, because the last intron in *talin* is small and the gene *CG6638* follows immediately after *talin*. All three exons were put back in the second step together with the sensor module resulting in one attR site in a *talin* intron and one in an *CG6638* intron. Respective controls with the individual fluorophores were also generated.

**Extended Data Fig. 2.**
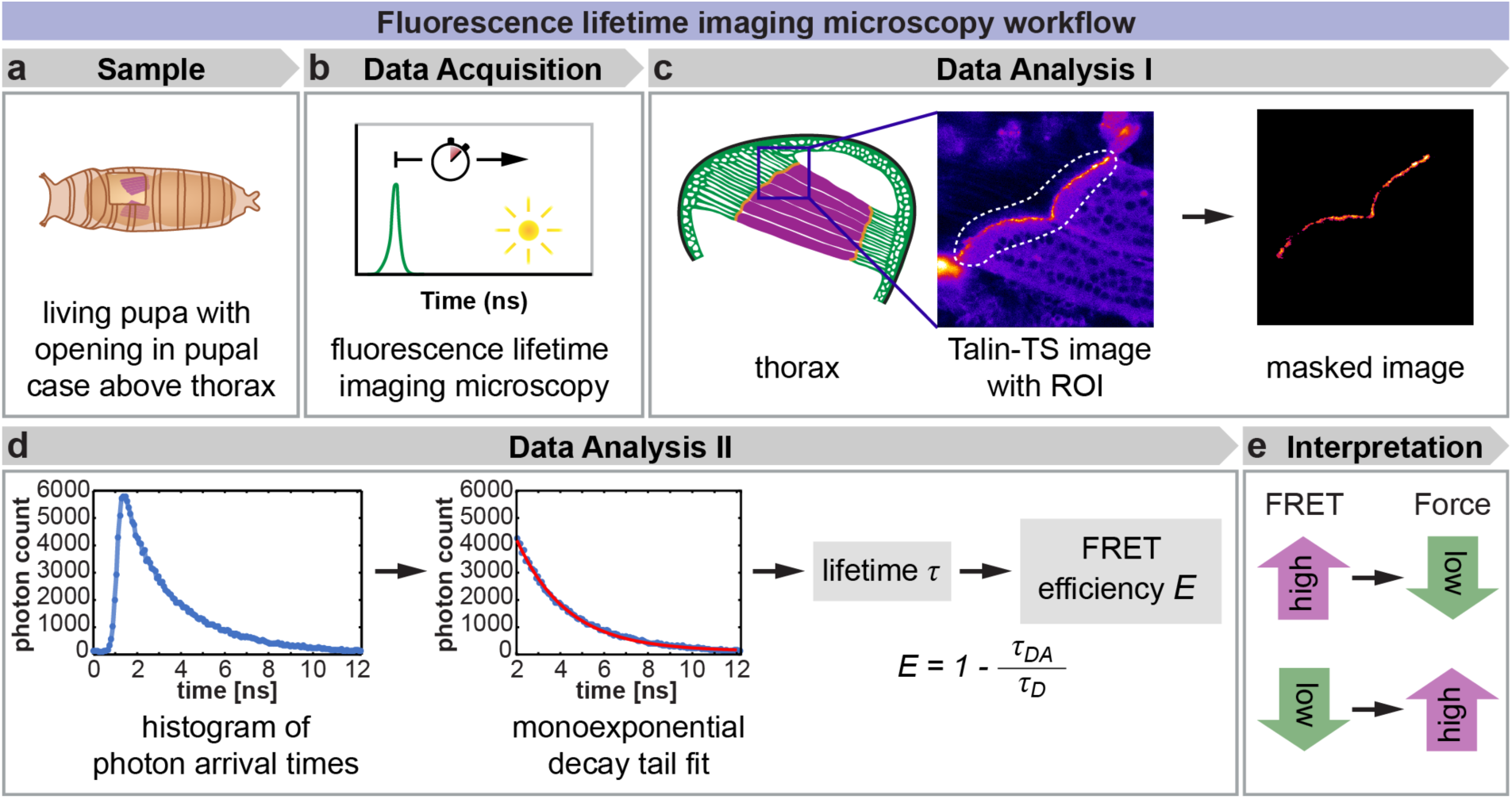
Fluorescence lifetime imaging microscopy workflow. **a**, Living *talin-TS* or control pupae were prepared for imaging by opening a window in the pupal case above the thorax containing the developing flight muscles (magenta)[50]. **b**, Fluorescence lifetime imaging microscopy (FLIM) was performed on a confocal microscope equipped with a pulsed laser (indicated by green peak) for exciting the donor fluorophore (YPet) and a time-correlated single photon counting (TCSPC)-detector for recording photon arrival times (indicated by yellow dot). **c**, A YPet intensity image created from the FLIM data was used to manually draw a region of interest (ROI) containing the anterior muscle attachments sites of the dorso-longitudinal flight muscles close to the surface of the thorax. From this ROI a mask for the muscle attachment sites was created by Multi-Otsu thresholding. **d**, Photon arrival times of all photons inside the mask were plotted in a histogram. The tail of the curve was fitted by a monoexponential decay to determine the lifetime *τ*. By comparing the lifetime of the Talin tension sensor *τ*_DA_ with the lifetime of respective donor-only control *τ*_D_, the FRET efficiency *E* was calculated. **e**, Interpretation of FRET results: A high FRET efficiency indicates mostly closed sensor modules and therefore low force. Vice versa, a low FRET efficiency indicates mostly open sensor modules and therefore high force.

**Extended Data Fig. 3.**
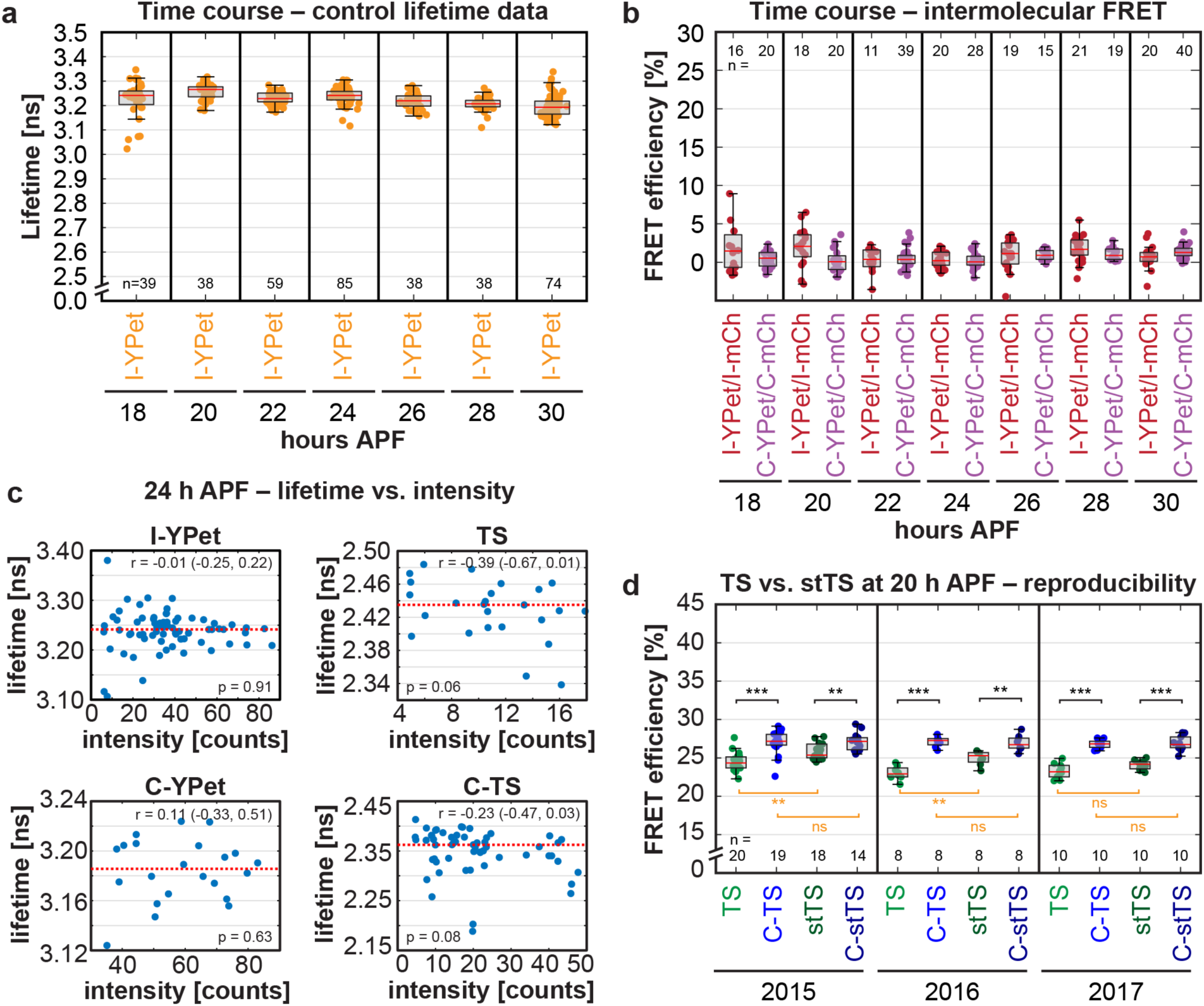
Control measurements for Talin forces detected at muscle attachment sites *in vivo*. **a**, Lifetime data of donor only controls at the internal position of Talin (I-YPet) **b**, Intermolecular FRET data measured by comparing heterozygous I-YPet/I-mCh or C-YPet/C-mCherry pupae to homozygous I-YPet or C-YPet pupae, respectively. Intermolecular FRET is negligible at all time points. **c**, Lifetime data for I-YPet, C-YPet, TS and C-TS at 24 h APF for each pupa plotted against the average intensity inside its muscle attachment site mask. Red dotted line represents median lifetime value. No correlation between lifetime and intensity could be detected (Pearson correlation coefficient r with 95% confidence interval and p-values are indicated). **d**, Reproducibility of FLIM-FRET measurements performed in different years: TS and its stable variant stTS show a reproducible decrease in FRET efficiency compared to the C-terminal zero-force controls C-TS and C-stTS at 20 h APF (Kolmogorov-Smirnov test, *** p<0.001, ** p*0.01, ns=not significant p>0.05; n=number of pupae).

**Extended Data Figure 4.**
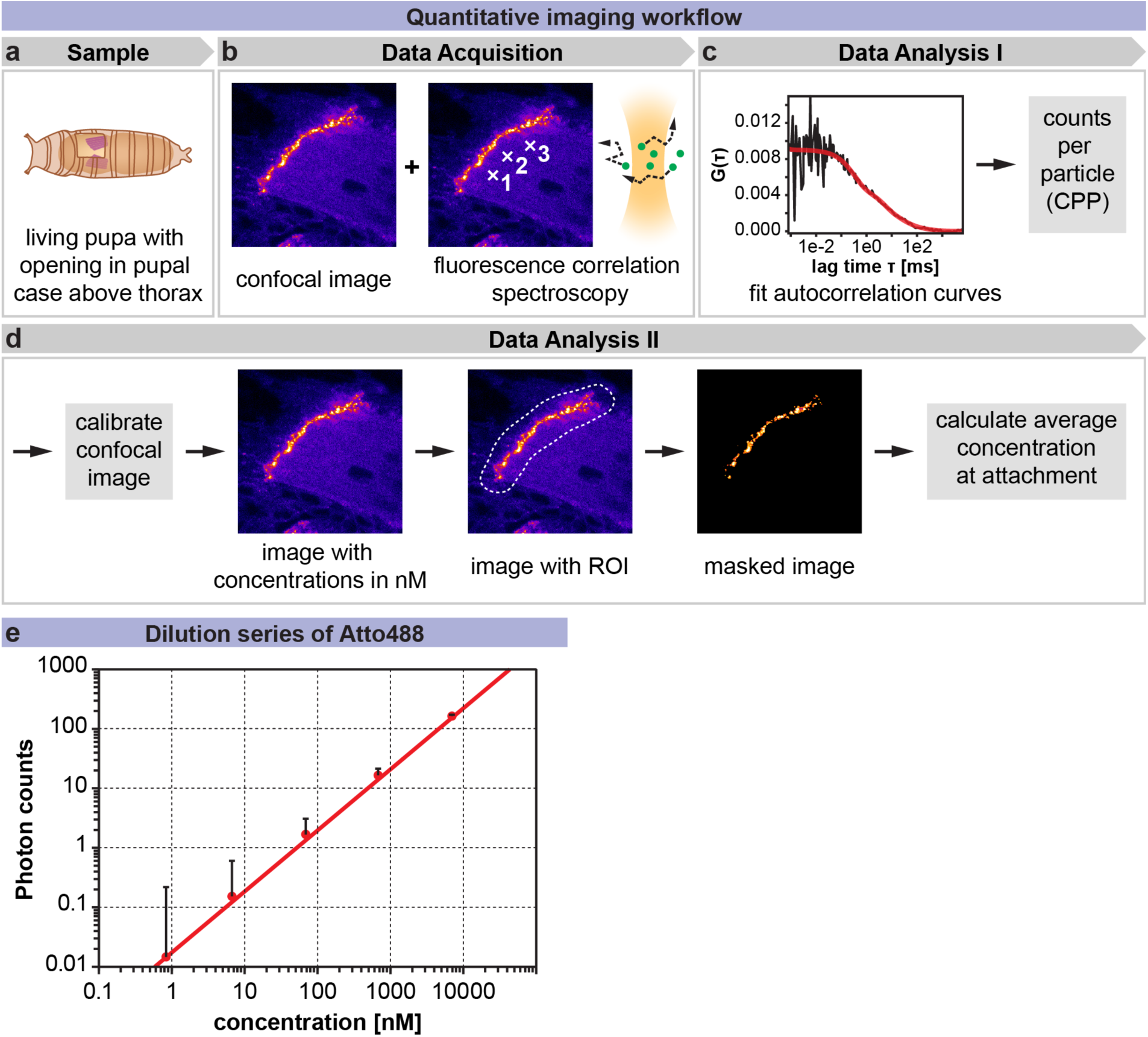
Quantitative imaging workflow and control measurements for fluorescence correlation spectroscopy (FCS). **a**, Living *talin-I-YPet* pupae were prepared for quantitative imaging by opening a window in the pupal case above the thorax containing the developing flight muscles (magenta)[50]. **b**, A confocal image and three FCS measurements were acquired using the same detector on a confocal microscope. **c**, Autocorrelation curves from the FCS measurements were fit to obtain a counts per particle (CPP) value for each pupa. **d**, The CPP value was used to calibrate each image resulting in a pixel-by-pixel concentration image. This image was used to manually draw an ROI around the muscle attachment site. From this ROI a muscle attachment mask was created automatically by Multi-Otsu thresholding. Finally, the average concentration at the attachment was calculated from the pixel-values inside the mask for each pupa. **e**, Pixel-by-pixel photon count values measured in an Atto488 dye dilution series (mean with standard deviation). Note that the photon count values increase linearly with the concentration of the dye for the entire range measured.

**Supplemental Video 1 – Legend**

Video of twitching primary muscle fiber shown in Fig. 2f. signal (green) is overlaid with the transmission light channel (grey) acquired simultaneously. The length of the movie is 1 min with a time resolution of 1.29 s played at 10x speed.

